# Disentangling effects of colocalizing genomic annotations to functionally prioritize non-coding variants within complex trait loci

**DOI:** 10.1101/009258

**Authors:** Gosia Trynka, Harm-Jan Westra, Kamil Slowikowski, Xinli Hu, Han Xu, Barbara E Stranger, Buhm Han, Soumya Raychaudhuri

**Affiliations:** Divisions of Genetics and Rheumatology, Department of Medicine, Brigham and Women's Hospital, Harvard Medical School, Boston, Massachusetts, USA; Partners Center for Personalized Genetic Medicine, Boston, Massachusetts, USA; Program in Medical and Population Genetics, Broad Institute of MIT and Harvard, Cambridge, Massachusetts, USA; Current address: Wellcome Trust Sanger Institute, Wellcome Trust Genome Campus, Cambridge, UK; Bioinformatics and Integrative Genomics, Harvard University, Cambridge, MA 02138, USA; Harvard-MIT Division of Health Sciences and Technology, Boston, MA USA; Department of Biostatistics and Computational Biology, Dana-Farber Cancer Institute and Harvard School of Public Health, Boston, Massachusetts, USA; Section of Genetic Medicine, Department of Medicine, University of Chicago, Chicago, Illinois, USA and Institute for Genomics and Systems Biology, University of Chicago, Chicago, Illinois, USA; Faculty of Medical and Human Sciences, University of Manchester, Manchester, UK

**Keywords:** complex traits, human disease, genetics, chromatin marks, ChIP-seq, DNase hypersensitivity sites, enrichment, fine-mapping

## Abstract

Identifying genomic annotations that differentiate causal from associated variants is critical to fine-map disease loci. While many studies have identified non-coding annotations overlapping disease variants, these annotations colocalize, complicating fine-mapping efforts. We demonstrate that conventional enrichment tests are inflated and cannot distinguish causal effects from colocalizing annotations. We developed a sensitive and specific statistical approach that is able to identify independent effects from colocalizing annotations. We first confirm that gene regulatory variants map to DNase-I hypersensitive sites (DHS) near transcription start sites. We then show that (1) 15-35% of causal variants within disease loci map to DHS independent of other annotations; (2) breast cancer and rheumatoid arthritis loci harbor potentially causal variants near the summits of histone marks rather than full peak bodies; and (3) variants associated with height are highly enriched for embryonic stem cell DHS sites. We highlight specific loci where we can most effectively prioritize causal variation.

---

Functional genomic annotations, including transcription factor binding sites and open chromatin regions, are rapidly becoming available^1–3^. These annotations provide valuable information for prioritizing potential causal variants within complex trait loci identified through genome-wide association studies (GWAS)^4–15^. However, the overlap of an associated variant with an annotation does not necessarily imply causality. While different functional genomic annotations might be most effective for prioritizing variants at individual loci, certain mechanisms might play a more dominant role for specific traits. So, annotations corresponding to those mechanisms may be able to prioritize causal variants for that trait. For example, binding sites for transcription factors regulating key pathogenic pathways might prioritize variants for diseases^8, 10^, or promoters active in a specific cell-type might be able to prioritize eQTLs variants from that cell-type.

Robust statistical strategies are required to identify informative annotations. A suitable strategy must control for two important types of potentially confounding genomic features: (1) local structure of genetic variation near SNP associations, and (2) colocalization of functional genomic annotations. First, trait-associated SNPs often map to regions with greater gene density, genetic variation, and more linkage disequilibrium (LD) compared to the rest of the genome. Second, functional annotations colocalize, and are often enriched within trait-associated loci. For example, DHSs colocalize with exons^16, 17^, and regulatory elements cluster together near and within gene transcripts. Therefore an observed enrichment for one annotation may be the consequence of unaccounted colocalization with other annotations, thus confounding inferences of causality.

Existing enrichment tests assess significance by comparison to matched SNP sets (**Supplementary Table 1**). These tests may be inflated if they fail to capture the genetic features at trait-associated loci. Therefore his we developed *GoShifter*, a statistical approach that accounts for these important confounders. To estimate the significance of overlap between trait-associated variants and annotations, we generate null distributions by randomly shifting annotations locally within a tested region. Additionally, we present a stratified test to identify independent contributions from colocalized annotations.

## RESULTS

### Summary of the statistical approach

To assess the statistical significance of an overlap between trait-associated SNPs and an annotation *X*, we first identify all variants in LD with each index SNP (r^2^>0.8 in 1000 Genomes Project^18^, Figure 1A). We then quantify the proportion of loci where at least one linked SNP overlaps *X*. To assess significance, we generate the null distribution by randomly shifting *X* sites within each trait-associated locus, and quantifying the proportion of overlapping loci (Figure 1B). We circularize each locus, so that shifted annotations remain within locus boundaries. This approach ensures that the null distribution maintains local genomic structures, such as locus size, number of SNPs in LD, gene density, and annotation density (**Online Methods**). We calculate a *delta-overlap* parameter that quantifies the difference between the observed overlap and the null. Larger delta-overlaps indicate larger proportions of overlapping variants, in a manner that is independent of the number of SNPs in LD and TSS or TES proximities of associated SNP sets (**Supplementary Figure 1**). For each locus, we also calculate an *overlap-score*, which reflects the probability of a specific SNP to overlap with *X* by chance.

**Figure 1.**
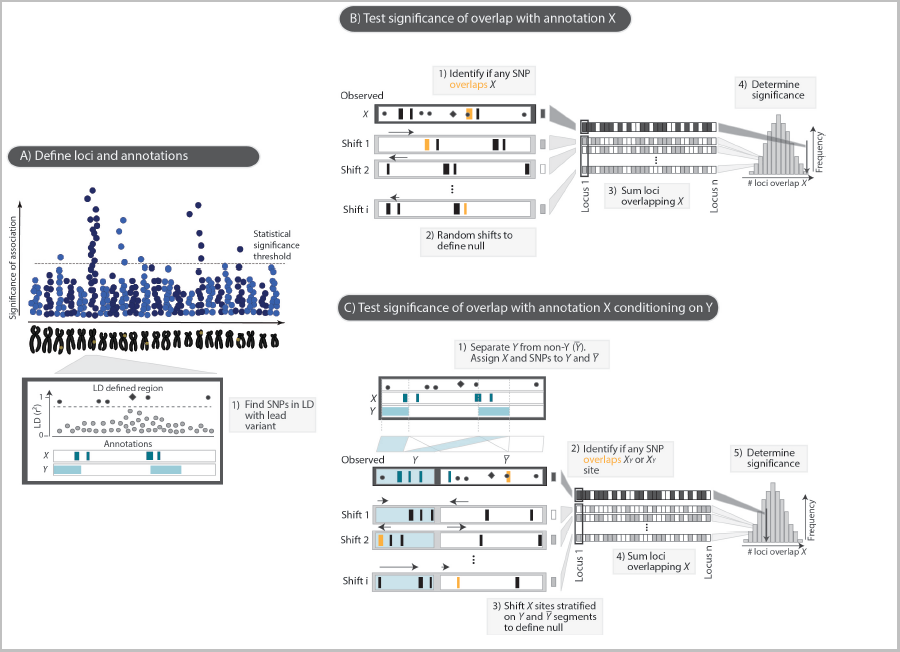
Schematic of the *GoShifter* method. To assess the statistical significance of an overlap between trait-associated SNPs and an annotation *X*, **(A)** we first use 1000 Genomes Project data to identify variants in LD with each index SNP (r^2^>0.8). **(B)** We quantify the observed overlap: the proportion of loci where at least one linked SNP overlaps annotation *X* (shaded boxes). The significance of observed overlap is estimated by comparing to a null distribution generated by randomly shifting *X* sites within each locus. After each shift, we calculate the proportion of loci overlapping the annotation. To ensure that the same number of shifted annotations remains within locus boundaries we circularize each region. **(C)** To determine the significance of an overlap with annotation *X* independent of a possibly colocalizing annotation *Y*, we partition each locus into two types of fragments: those regions mapped by *Y* sites (light blue blocks), and those that lack them (denoted as *Ȳ*; white blocks). We join the respective *Y* and *Ȳ* fragments into two independent continuous segments. To generate the null distribution, we shift annotation *X* separately within each of the two segments. For each iteration we count the proportion of loci where any of the linked SNPs overlaps annotation *X* in either *Y* or *Ȳ*.

To assess the significance of overlap of trait-associated SNPs with annotation *X*, while accounting for a second (possibly colocalizing) annotation *Y* we present a stratified approach. To do this, we partition each locus into two subsegments based on the presence of *Y*: (1) the set of *Y* annotated regions (denoted as *Y)*, and (2) the set of regions lacking *Y* (denoted as *Ȳ*; Figure 1C). We then merge the *Y* regions together and the *Ȳ* regions together for each locus into separate subsegments. Ultimately each merged subsegment includes variants and *X* annotations. Contiguous *X* annotated regions may be split if only parts of it are overlapping *Y*. This approach stratifies each locus, fixing the overlap of the SNPs to *Y*, as well as the spatial relationship between *X* and *Y*. We then generate the null distribution by shifting *X* sites within the *Y* and *Ȳ* subsegments separately, and then quantifying the proportion of loci in which any linked SNPs overlaps *X* in either *Y* or *Ȳ*. If *X* contains information independent of the colocalization with *Y*, stratified shifting should reduce the number of loci where linked SNPs randomly overlap *X*.We implemented both unstratified and stratified analyses in our method, called *GoShifter*.

### GoShifter demonstrated appropriate statistical properties; standard enrichment tests demonstrated inflated statistics

To evaluate the statistical properties of *GoShifter* and other enrichment tests, we simulated 1,000 sets of 1,416 trait-associated SNPs resembling variant associations identified through GWAS (**Online Methods**). Unlike real trait-associated variants where the function and identity of causal variants is unknown, we designed our SNP sets based on causal variants that mapped to specific genomic annotations (**Supplementary Figure 2**), such as DHS, promoters, 5'UTRs, exons, 3' UTRs, introns, and intergenic regions. We tested these 1,000 SNP sets for overlap with DHS regions consolidated from 217 cell types (**Online Methods**). Choosing causal SNPs to overlap specific annotations allowed us to assess a method's ability to identify true enrichment and reject spurious overlap (**Supplementary Figure 3**). An appropriate strategy should detect DHS enrichment in SNP sets that were designed to tag functional variants in regulatory regions (DHS, promoters, 5'UTRs), modest enrichment for exons and 3'UTRs (which colocalize with DHS)^16, 17^, but not at introns or intergenic regions (**Online Methods**).

*GoShifter* was well powered to detect significant enrichment in simulated SNP sets that tagged variants in DHS, promoter, and 5'UTR regions; 100% of such SNP sets obtained p<0.001 based on 1,000 shifting iterations (Figure 2A and **Supplementary Figure 4**). By chance, we would expect 5% of the SNP sets tagging variants in intronic and intergenic regions to obtain p<0.05. Indeed, we observed that 4.44% and 7.4%, respectively, obtained p<0.05 (Figure 2A, **Supplementary Table 2**). The SNP sets tagging exons and 3'UTR appropriately showed modest enrichment (60% and 11% of these SNP sets obtained p<0.05, respectively).

**Figure 2.**
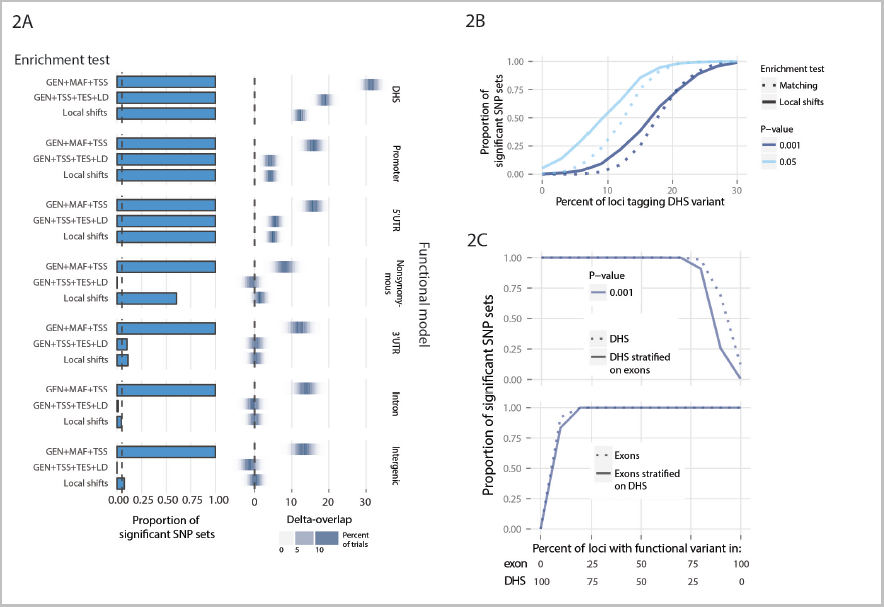
Comparison of statistics between GoShifter and matching-based tests. **(A)** The most widely used matching-based test (GEN+MAF+TSS) results in highly significant DHS overlap even in the SNP sets that do not tag regulatory elements (e.g. introns). This is corrected by including LD and TES parameters in matching. However, even though this improves the false positive rate in intergenic SNP sets, this test is inappropriate in more complex SNP sets, such as non-synonymous variants, where it fails to recognize true enrichment. **(B)** *GoShifter* is better powered than stringent matching to detect DHS enrichment in SNP sets with 0 to 30% of loci tagging DHS variant at p<0.05 and p<0.001. **(C)** Without accounting for possible colocalization between exons and DHS, 12.5% of SNP sets show significant DHS enrichment when all loci tag exonic variant (at p<0.001). This drops to 0.7% when testing for DHS enrichment and stratifying on exons (upper panel). *GoShifter* retains the power to detect enrichment for exons across SNP sets with 10% loci tagging exon variants, even when stratifying on possible colocalization with DHS (lower panel).

We also tested commonly used matching-based enrichment tests^1, 5,8,10–13,15,19–28^ (**Supplementary Table 1**). These tests assess significance by comparing observed overlap to that of SNP sets randomly sampled from the genome. Typically, such methods match SNPs based on localization within or outside a gene (GEN), minor allele frequency (MAF), and proximity to transcription start site (TSS)^1,5,10,15,20,24,27^. All simulated SNP sets, including 100% of those deriving causal variants from intergenic regions, obtained p<0.05 for DHS enrichment (Figure 2A, **Supplementary Table 2**), using any matching-based enrichment tests. This demonstrates that matching strategies can result in highly inflated statistics that might produce false positive results.

We further tested the effectiveness of matching on different combinations of genomic features, and observed that the results were highly sensitive to the choice of specific matching parameters. We noted that the number of SNPs in LD is not typically used as a parameter, yet crucial for reducing inflated statistics (**Supplementary Figure 4)**. While MAF was frequently included, it has little effect. Matching on the combination of GEN, TSS, TES, and LD adequately controlled type I error, if the SNPs were selected from non-regulatory regions (e.g. intergenic or intronic regions).

### GoShifter outperformed stringent SNP-matching

We calculated the power to detect DHS enrichment, using SNP sets harboring various proportions of functional variants within DHS. *GoShifter* outperformed SNP-matching on GEN, TSS, TES and LD (Figure 2B). When 10% of a SNP set tagged DHS, *GoShifter* had a 55% power to detect enrichment, compared to 31% with stringent SNP-matching (Figure 2B).

### Differentiating effects of colocalizing annotations

To test the ability of *GoShifter* to control for the effects of colocalized annotations, we examined two scenarios. First, we queried the enrichment of different annotations in 1,000 sets of 1,416 SNPs tagging variants selected from exon sites. While we observed appropriate enrichment with exons across all SNP sets (p<0.05), we also saw DHS enrichment in 60% of them (Figure 2A). This secondary enrichment was a result of colocalization between the DHS sites and exons^16,17^. Testing for DHS enrichment after stratifying on exons produced much lower type I error rate with only 10.2% of SNP sets obtaining p<0.05. The modest residual inflation was likely because our simulation strategy used array-based SNPs (Online Methods), which are enriched for DHS^5^.

Second, we assessed 1,000 sets of SNPs tagging variants within DHS. While all SNP sets were enriched for DHS at p<0.05, 98% of the SNP sets were also enriched for H3K4me3 sites, which highlights promoters and colocalizes with DHS. Since H3K4me3 enrichment was likely secondary to functional variants in DHS, we expected the effect to become insignificant once we stratified on DHS. After stratifying on DHS, only 1.3% of the SNP sets showed H3K4me3 enrichment at p<0.05, confirming that it was a non-independent effect. Conversely, when tested for enrichment stratifying on H3K4me3, DHS overlap remained significant in 100% of SNP sets.

The stratified analysis only modestly compromises power to detect effects. For SNP sets where causal variants were generated from DHS and exons, stratifying on exons reduced power to detect DHS enrichment at p<0.001 only when ≤ 20% of the causal variants were from DHS (Figure 2C). However, stratifying on DHS sites did not limit the power to detect exon enrichment at all.

### eQTL SNPs are enriched for DHS and enhancers close to transcription start sites

To demonstrate the applicability of our approach with real data, we aimed to identify causal genomic annotations that overlapped 6,380 expression quantitative trait loci (eQTLs) that influenced gene expression. We selected eQTLs as an instructive example since they are known to localize near the TSS and sites involved in active gene regulation^29,30^. We observed that the TSS proximity (+/−0.5kb, 1kb, 2kb, 5, 10kb, 25kb and 50kb) and DHS consolidated from 217 cell types each showed highly significant enrichment (p<10^−4^; Figure 3A). We tested the DHS overlap while stratifying on different TSS proximity windows, and observed highly significant independent enrichment (p<10^−4^). In contrast, when we tested TSS proximity stratifying on DHS, we detected significant enrichment only at sites very close to the TSS (+/−500 bp, p=2×10^−4^). Furthermore, DHS located within 5kb around the TSS demonstrated the strongest signal. Genome-wide DHS sites were not significantly associated when we stratified on DHS sites 10kb around the TSS region (p>0.03). These results suggested that causal variants for eQTL-SNPs influence gene expression largely (but not exclusively) through specific open chromatin sites in close proximity to TSS.

**Figure 3.**
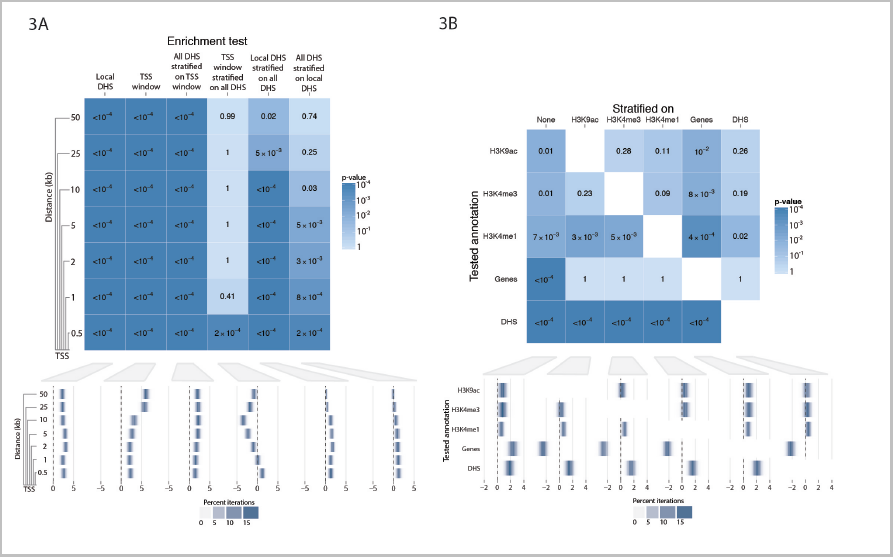
eQTL variants localize to DHS near TSS regions. **(A)** Results and the effect size of the enrichment for DHS and proximity to TSS across 6,380 eQTL variants. We examine windows around the TSS and assess significance of eQTL overlap. **(B)** Enrichment result across active regulatory chromatin marks and gene transcripts with eQTL SNPs. Upon stratifying on DHS, association of different annotations is reduced. This is particularly apparent for gene transcripts, indicating that the observed enrichment is due to the colocalization with gene regulatory elements.

We also observed enrichment (p<0.05) for histone modifications that indicated active gene regulation (H3K9ac, H3K4me3 and H3K4me1) as well as for gene transcripts (Figure 3B). Stratified testing on combinations of all annotations immediately revealed that the observed enrichment for transcripts was due only to colocalization with DHS. DHS showed the strongest enrichment and remained significant when stratified on all other annotations (p<10^−4^). We also observed a nominal independent enrichment at H3K4me1 sites (p=0.02). Therefore, causal eQTL variants might be best prioritized by identifying SNPs (1) in DHS sites, (2) within close proximity to the TSS (up to 5kb) or (3) within H3K4me1 regions.

### Quantifying the proportion of GWAS catalog SNPs with DHS causal variants

We assessed 1,416 independent SNP associations from the NHGRI GWAS Catalog^31^ for overlap with different annotations (**Online Methods**). We observed enrichment at DHS (p<10^−4^), and evidence of enrichment at genes, H3K4me3 and H3K4me1 marks, as well as 5 and 10kb windows around TSS (Figure 4A). Pairwise stratified tests showed that DHS sites were independently enriched from other annotations (p≤7×10^−4^). In contrast, the enrichment for gene transcripts and TSS regions were not significant after stratifying on DHS. Both H3K4me1 and H3K4me3 retained nominal enrichment independent from each other and DHS (p≤0.05; Figure 4B). These results suggest that causal disease-associated variants in DHS sites function through mechanisms independent of those indicated by H3K4me3 and H3K3me1.

**Figure 4.**
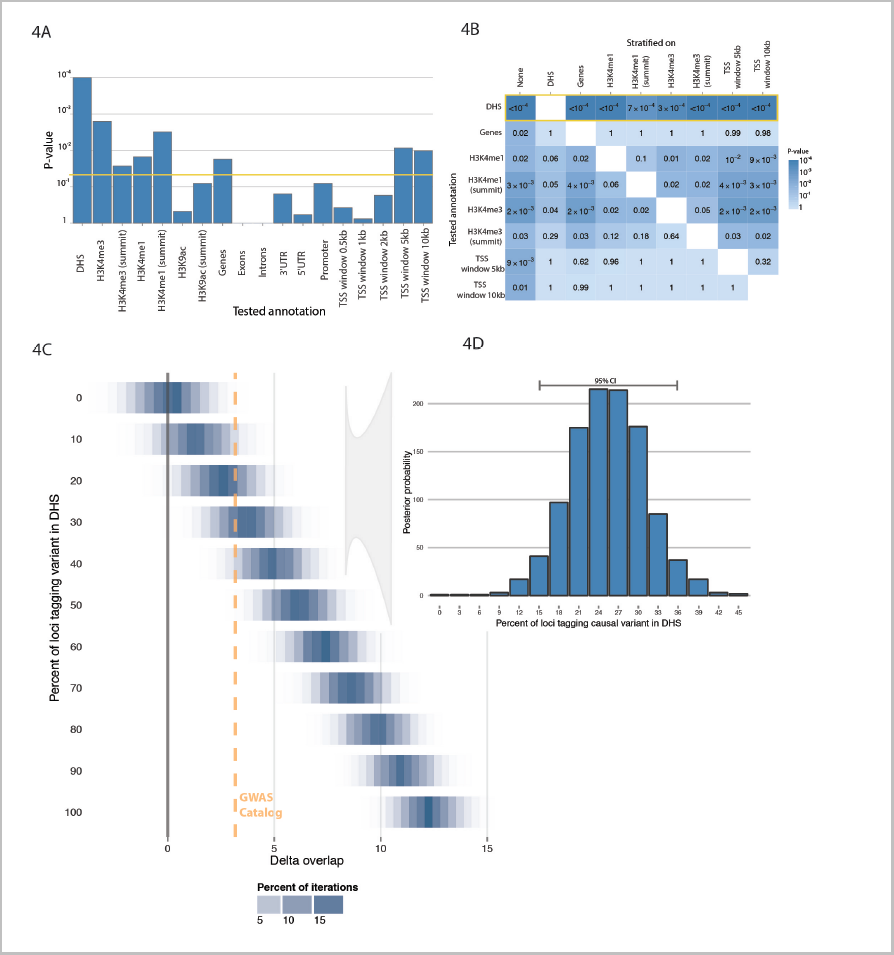
Quantifying the proportion of causal variants within the GWAS catalog derived from DHS. **(A)** The enrichment results of different genomic annotations with 1,416 independent NHGRI GWAS Catalog SNPs. **(B)** Pairwise stratified analysis implicates that the primary driver of the enrichment is DHS. Additionally, H3K4me3 and H3K4me1 summit regions are independently enriched but to a lesser extent than DHS. **(C)** We estimate delta-overlap for DHS at 3.17, **(D)** which corresponds to 15-36% of loci with causal variants within DHS.

Previous reports estimated that 40-80% of trait-associated variants within the GWAS catalog overlap with DHSs^1,5,15^. To accurately determine the proportion of GWAS loci that tag variants in DHS, we compared the delta-overlap, which quantifies the strength of association, for DHS observed in the GWAS catalog with delta-overlap values for DHS sites using 11,000 sets of 1,416 randomly selected independent SNPs with variable proportions of causal SNPs in DHS sites (**Online Methods**; Figure 4C). For DHS overlapping the GWAS catalog, we observed a delta-overlap of 3.17. Delta-overlap is independent of the number of SNPs in LD and TSS or TES proximities of associated SNP sets (**Supplementary Figure 1**). We determined that this value was consistent with having 15-36% causal variants within DHS (95% confidence, Figure 4D), suggesting DHS enrichment may be more modest than previously reported.

### Rheumatoid arthritis and breast cancer associations are enriched at the summits of cell-type specific histone marks

We examined two phenotypes with large numbers (>50) of associated variants, to test GoShifter's ability to identify cell-type specific functional variants. We first tested 88 SNPs associated with rheumatoid arthritis (RA)^32^ for enrichment of non-coding annotations. We focused on CD4^+^ memory T cells given recent observations of cell-type specific gene expression and eQTL within these loci^4,33,34^. We observed no association with H3K4me3 peak bodies in CD4+ memory T cells (p=0.17) or in the aggregate of all 118 cell types from our datasets (p=0.14, Figure 5A). Since the median width of H3K4me3 mark peaks varied widely (110bp to 86,490bp), we examined the summit regions (+/−100bp of H3K4me3 summits) where active gene regulatory elements were most likely located. In the summit regions we observed significant overlap both for the 118 cell types collectively (p=0.044) and CD4+ memory T cells specifically (p=1.6×10^−3^). The CD4+ memory T cell signal remained significant after stratification on the summit regions of the other 117 cell types (p=2.7×10^−3^). In contrast, the other cell types did not retain significant enrichment after stratifying on CD4+ memory T cells (p=0.08). These results suggested that examining the H3K4me3 summit regions in CD4+ memory T cells could help prioritizing causal variants in RA loci.

**Figure 5.**
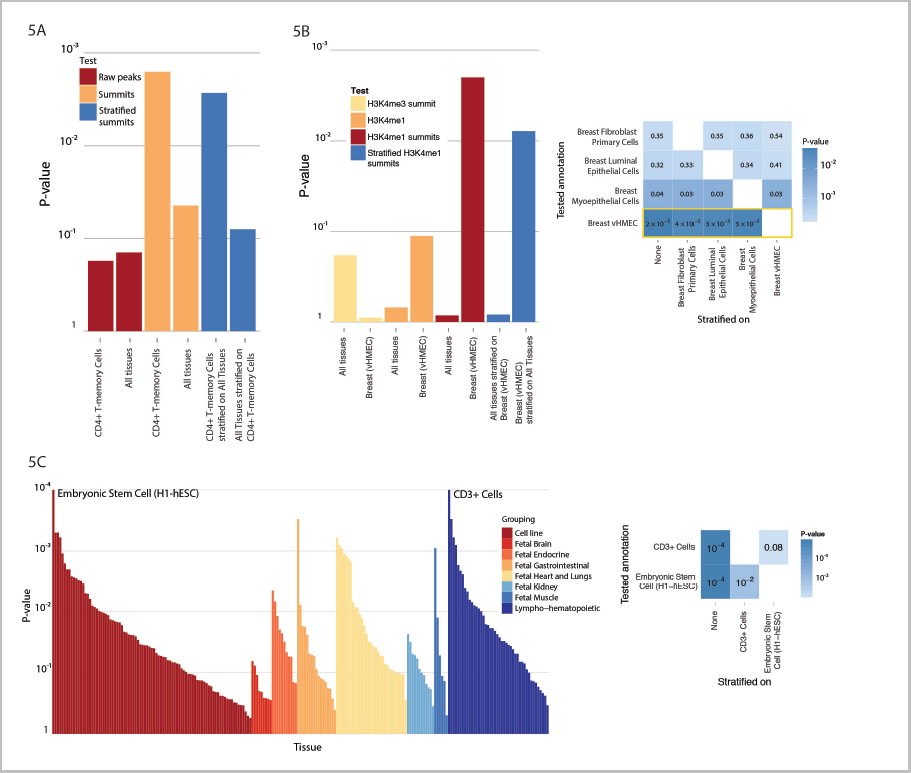
Enrichment results for three selected sets of trait-associated SNPs. **(A)** Rheumatoid arthritis variants show highly significant enrichment within summit regions of H3K4me3 peaks from CD4^+^ memory T cells. **(B)** Breast cancer SNPs are highly enriched for summit regions of H3K4me1 peaks in variant human mammary epithelial cells (vHMEC; p=2×10^−3^; left panel). This enrichment is maintained when stratifying on summit regions from each of the other three breast tissues (p<3.6×10^−3^; right panel). **(C)** SNPs associated with height show high enrichment for DHS regions in embryonic stem cells and CD^3+^ cells from cord blood (left panel). However, the CD^3+^ cell DHS enrichment diminishes after stratifying on embryonic stem cells (p=0.08), while embryonic stem cells retain significance when stratifying on CD^3+^ cells (p=9.6×10^−3^; right panel).

Similarly, we examined 69 SNPs^35^ associated with breast cancer. H3K4me3 summit regions revealed no significant overlap with the three breast tissues present in our dataset (p>0.4). We therefore tested for enrichment with another active regulatory mark, H3K4me1, for which there were four breast tissues in our dataset. This mark identified only breast myoepithelial cells with nominal significance (p=0.034). However, when we used H3K4me1 summit regions we observed more significant overlap for variant human mammary epithelial cells (vHMEC; p=2×10^−3^; Figure 5B). We performed pairwise stratified enrichment tests across the four breast tissues, and found that the vHMEC summit region enrichment was maintained when stratifying on peaks from each of the other three breast tissues (p<3.6×10^−3^). DHS appeared less informative for breast cancer, as we found that none of the DHS samples in our dataset showed nominally significant enrichment. These results show that our stratified approach can be applied to narrow down the specific cell type in which disease associated variants function, even from within the disease tissue of interest.

### Stratified analysis can indicate relevant cell types for height

We assessed 697 SNPs associated with adult human height^36^, a highly polygenic trait without clearly established causal cell types. When we examined DHS from 217 cell types collectively, we observed nominal evidence of overlap (p=0.019). Individually, many tissues demonstrated some evidence of overlap, including 13 at p<10^−3^ (Figure 5C). We observed the strongest enrichment in embryonic stem cells (H1-hESC; p<10^−4^) and primary CD3+ cells from cord blood (p<10^−4^). The enrichment in H1-hESC retained significance after stratifying on cord blood CD3+ cells (p=9.6×10^−3^); but the CD3+ cell DHS enrichment diminished after stratifying on H1-hESC (p=0.08). Examining DHS in embryonic stem cells or a related cell type might be informative for fine-mapping height loci for potential causal variants.

### Fine-mapping to functional variants

Once critical annotations had been identified, we used the *overlap-score* from *GoShifter* to identify loci where variants might be best prioritized. In breast cancer, the lowest *overlap-score* was the rs889312 locus (score=0.097, Figure 6). This SNP was in LD with seven other variants, of which only rs1862626 overlapped a vHMEC H3K4me1 summit region. This SNP is upstream of *MAP3K1* and modifies a predicted binding site for estrogen receptor alpha (ER-α)^37^, supporting the well-established role of estrogen-mediated signaling in breast cancer progression^38,39^. For height, the rs11677466 locus showed the lowest *overlap-score* (0.026; Figure 6). This SNP itself overlaps an embryonic stem cell DHS site that is a known binding site for the HNF-4α^40–42^, a transcription factor that plays important roles in metabolic regulation and stem cell differentiation.

**Figure 6.**
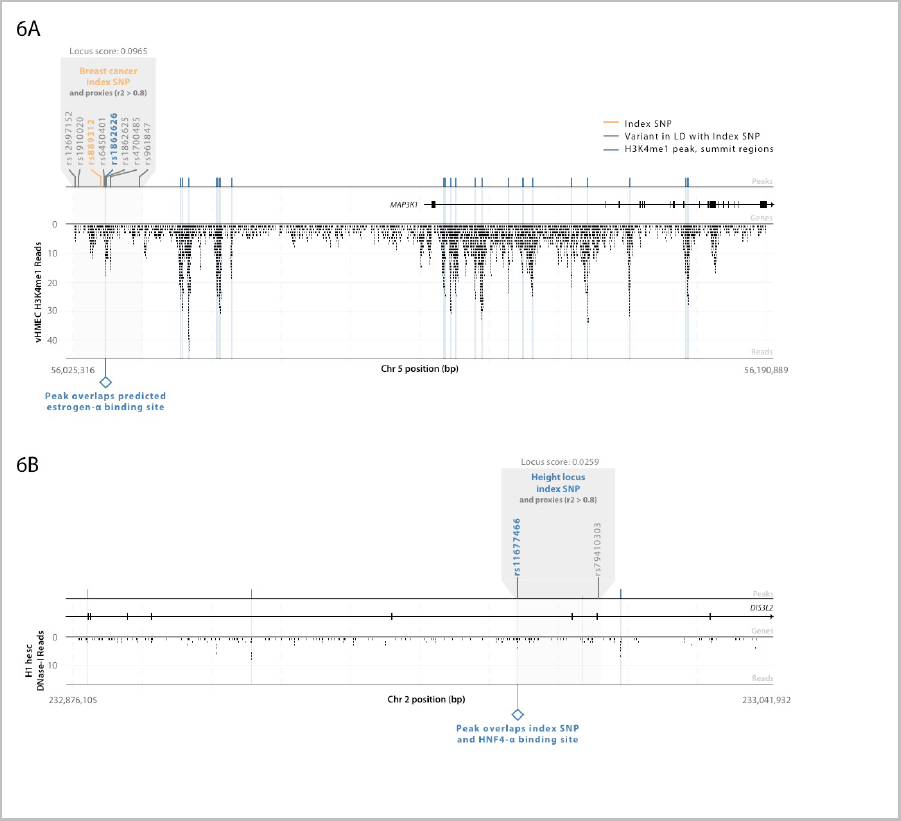
Locusplots showing the peaks, variants and reads in two trait-associated loci. **(A)** The locus with the best overlap-score among breast cancer SNP associations was the locus marked by the index SNP rs889312. This SNP has a variant in LD (r^2^>0.8; rs1862626) that overlaps with the summit region of an H3K4me1 peak. This peak overlaps with a predicted estrogen-alpha binding site. The associated locus is located upstream of potential oncogene *MAP3K1*. **(B)** Of the the height assocated SNPs, the locus with the best overlap-score was the one marked by the index SNP rs11677466, located in an exon of *DIS3L2*. This SNP overlaps a DNase-I hypersensitivity peak, which also overlaps a known HNF4-alpha binding site.

## DISCUSSION

To date, most published studies estimate the statistical significance of the overlap of associated variants with genomic annotations by using a random sets of matched SNPs for comparison. These approaches are vulnerable to inflated statistics and false discoveries, and might overestimate the utility of certain genomic annotations (see **Supplementary Figure 5**). For instance, we estimate that DHS accounts for 15-36% of GWAS catalog variants rather than the widely reported 40-80%. We note that matching-based enrichment tests for DHS must at least include TSS, TES, LD, and GEN as parameters. Other annotations may require different parameters sets, adding additional challenges to using such approaches.

*GoShifter* stringently controls for local features by shifting annotations within trait-associated loci. Similar approaches have been employed to study the relationship of variants with genomic annotations and different annotations with each other^4,27,43,44^. In addition, our method represents an important advance over current approaches in its ability to assess independent effects from colocalizing annotations. One potential limitation of this approach is that it cannot easily account for the biases intrinsic to commercial arrays; however as studies increasingly use whole-genome imputation and sequencing, this limitation will become less relevant.

Applying *GoShifter*, we observed that different annotations play dominant roles in different traits. For example, height-associated variants showed DHS enrichment specifically in embryonic stem cells, while breast cancer variants enriched for H3K4me1 but not DHS. These may be partly due to different trait-relevant tissues being mapped with different assays. Nevertheless, variants associated with different phenotypes are likely to act through various mechanisms, resulting in differential enrichment of annotations.

Once the most informative annotations and trait-relevant tissues are identified, it becomes possible to functionally fine-map trait-associated regions. For example, we demonstrated that the summit regions of histone marks are more informative than the entirety of histone mark sites for breast cancer and rheumatoid arthritis. We anticipate that the applications of accurate enrichment statistics will only increase as more loci are discovered and better localized with dense-genotyping and sequencing, and as the diversity and quality of genomic annotations expand.

## ACKNOWEDGEMENTS

We acknowledge the GIANT consortium for providing us access to the height SNPs. We also acknowledge Joel Hirschhorn, *X* Shirley Liu, Mark Daly, Tonu Esko, Dorothee Diogo, Joshua Randall, for helpful scientific discussions. This work is supported by funding from the National Institutes of Health (NIAMS-1R01AR063759 and NHGRI-1U01HG0070033 to SR), a Doris Duke Clinical Scientist Development Award (SR), and the Rubicon grant from The Netherlands Organization for Scientific Research (GT).

## AUTHOR CONTRIBUTIONS

S.R. led the study. G.T. performed the analysis. H.X. provided the DHS data. G.T., H-J.W., X.H., B.H., K.S., B.E.S and S.R. wrote the manuscript. All authors reviewed the final manuscript.

## ONLINE METHODS

### Enrichment Tests: Assessing the Significance of Overlap of Variants and Functional Genomic Annotations

Typically, *enrichment tests* first quantify the proportion of loci that overlap with a tested annotation, and then assess if the observed overlap exceeds chance using a null distribution of expected overlap. This null distribution can be derived by sampling random sets of SNPs from the genome with properties that match the tested SNPs, or by locally shifting the annotations within the tested loci, as we propose here.

### A. Assessing enrichment by local annotation shifting.

One approach is to define the null distribution of overlap by locally shifting annotations within loci^4,27,43,44^, which we implemented in *GoShifter* (see Figure 1). Shifting annotations locally ensures that we always test the same number of annotations and the same number of linked SNPs. Also, it preserves the structure of the annotations within each locus.

We first determined the median size of the tested annotation. We then defined the locus boundaries based on the position of the most upstream and downstream SNPs in LD (r^2^ > 0.8), and further extended by twice the median size of the tested annotation. This ensured that variants without SNPs in LD had non-zero locus sizes. We identified all annotation sites within each locus boundaries, and determined the number of loci where at least one SNP overlapped with the annotation. To construct the null distribution of chance overlap, we circularized the region, which prevents the annotation from being shifted outside the locus boundaries. We then fixed the positions of the SNPs in the locus while shifting all annotations by a random distance, preserving the distances between annotation intervals. For each shift, each locus is assigned a random shift value ranging from −*l_i_*; to +*l_i_*, where *l_i_* represents the size of the locus *i*. At each iteration, we count the number of loci with at least one SNP overlapping the annotation after shifting, which estimates the number of loci that overlap by chance. Over all iterations, we create a null distribution from which we compute the p-value as the proportion of iterations for which the number of overlapping loci is equal to or greater than that for the tested SNPs. For all analyses in this paper that used simulated SNP sets, we conducted *n* = 10^3^ shifts. To test for enrichment of trait-associated variants we performed *n* = 10^4^ shifts.

### Delta-overlap parameter.

To quantify the effect size of the observed enrichment we introduce the *delta-overlap* parameter. We define the proportion of loci overlapping the annotation under the null by shifting as described above. The *delta-overlap* is the difference between the observed proportion of loci with overlap and the mean of the null distribution. If there is no enrichment, the observed real overlap will be close to the mean overlap observed under the null, and *delta-overlap* will be close to zero. Conversely, larger *delta-overlap* values correspond to stronger enrichment.

### Overlap-score parameter

We define the *overlap-score* as the likelihood of each locus to overlap an annotation beyond the overlap expected by chance. This is computed only for the loci that overlap the annotation in the observed data. *Overlap-score* is defined as 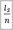 where *I_s_* is the number of shifting iterations where at least one SNP overlaps the annotation and *n* is the total number of iterations. A small score reflects a low probability that the overlap will happen by chance. The loci with low scores are the ones likely to be driving significant observations, making them higher-priority candidates for further functional investigations.

### B. Stratified enrichment of an annotation, controlling for the effect of a secondary annotation.

We further extend the shifting enrichment test to assess the significance of overlap of an annotation *X*, controlling for any overlap with a potentially colocalizing annotation *Y* (see Figure 1). This approach allows the comparison of multiple annotations against each other. To do so, we divide each locus into regions overlapping *Y* and regions not overlapping *Y*, and treat them independently in a “stratified” analysis.

For each associated locus we map all annotations *X* (to be tested) and *Y* (to be conditioned on) that fall between region boundaries. We first identify segments within the locus into fragments that either overlap *Y* or not. We fuse all *Y*-overlapping fragments into one continuous segment (denoted *Y*), and similarly fuse all non-Y-overlapping fragments into another segment (denoted as *Ȳ*). When we fuse the segments we include overlapping variants and regions annotated with *X*. If an *X* annotation does not fall entirely within *Y* or *Ȳ*, it is split at the boundary into the *Y* and *Ȳ* segments. As the result, the relationships and relative positions between *X*, *Y*, and the SNPs in the locus are preserved in both segments.

We then count the number of loci within which at least one SNP overlaps with annotation *X* in the *Y* or *Ȳ* segment. To evaluate the statistical significance of the observed enrichment, we shift annotation *X* within segment *Y* and *Ȳ* independently. We circularize each of the two segments *Y* and *Ȳ* to ensure that an annotation cannot fall outside the segment boundaries. At each iteration, we sum the number of loci containing at least one SNP that overlaps annotation *X* in one of the segments. We define the enrichment p-value as the proportion of iterations where the number of loci with SNP overlapping *X* exceeds the number of loci overlapping *X* prior to shifting

### C. Assessing enrichment by SNP matching.

The common strategy to assess enrichment of an annotation is to compare associated SNPs to random matched SNPs in an attempt to control for possible genomic confounders, however there is no consensus as to which parameters are relevant for the matching-based tests (**Supplementary Table 1**).

We noted that different genotyping platforms had distinct biases due to their designs [ref] (efficiency of tagging, allele frequency of included SNPs, number of SNPs, and physical distribution in the genome). To avoid such bias produced by platform differences, here we derived the null distribution by sampling variants from the same platform that we used for simulating the SNP sets (Illumina Omni 2.5).

To match the set of SNPs being tested for enrichment, we binned them by GEN (gene overlap), MAF (frequency bins incremented by 5%), TSS proximity (defined by 500bp, 2kb, 5kb, 10kb, 20kb and 100kb distance from the nearest TSS), TES proximity (defined by 1kb, 2kb, 5kb, 10kb, 20kb and 100kb distance from TES of the same gene as the nearest TSS), and the number of SNPs in LD. If fewer than 20 SNPs were present in the matched bin, we expanded to the nearest LD bins while matching on the other properties. We constructed the null distribution by repeatedly sampling the overlap between the annotation and the matched SNP sets. Then, we calculated the enrichment p-value as the proportion of matched SNP sets with a number of overlaps equal to or greater than the tested SNPs.

### Genomic Annotations

Our study utilized several commonly used functional genomic annotations, as well as gene annotations. These annotations were used to test enrichment for different variant sets, and in some cases were used to generate simulated SNP sets to assess statistical properties. All of these annotations were compiled from publically available resources.

### DNase Hypersensitivity Data

We used the DHS data from 80 experiments from ENCODE^1^ and 137 from NIH Roadmap Epigenomics Project^2^. See **Supplementary Table 3** for a detailed list of included tissues and cell types. We consolidated DHS tracks across all cell-types for analyses on eQTL and GWAS catalog data (as in **Figures 2, 3, 4**), but for individual cell-types for height data (as in Figure 5C and 6). For both datasets we downloaded hg19-mapped ChIP-Seq reads. We merged reads from samples with multiple replicates into a single library. For each sample, we used the corresponding input DNA library as the control if available. We ran MACS software (v2.0)^45^ using default settings (FDR= 0.01 and bandwidth = 300 bp) to identify significant peaks by comparing the DHS libraries with the control. For each experiment we located the start and end positions of the peaks to define genomic intervals in a BED file. We combined replicates so that any overlapping DHS peaks from different replicates were merged into a single interval. In total, the 217 experiments consisted of 1,331,772 distinct autosomal DHS positions, collectively spanning 16.4% of the genome.

### Histone Modifications

Similarly, we used MACS software (v2.0) with default parameters to call H3K4me3, H3K4me1 and H3K9ac peaks in 118, 114, and 50 tissue and cell type samples, respectively, from the NIH Roadmap Epigenome^2^. We examined histone mark data by aggregating across all cell-types for eQTL and GWAS catalog data, and for individual cell-types for breast cancer and rheumatoid arthritis. See **Supplementary Table 3** for a detailed list of included tissues and cell types. We downloaded data from http://www.genboree.org/EdaccData/Current-Release/experiment-sample/ as available on Jan 2, 2014. If multiple replicates of the same tissue (input and control) were available, we provided multiple BED files as input for MACS. For each experiment we located the start and end of the peaks, as well as the summit regions defined as +/−100bp around summits called by MACS.

### Gene Annotations

We defined gene annotations, including gene (whole transcript), exons, 5' UTR, 3' UTR, intron and promoter regions. To this end, we downloaded the RefSeq gene coordinates from the UCSC Genome Browser (http://genome.ucsc.edu/). We retained only transcripts with more than one exon. To exclude poorly studied genes, pseudogenes, and falsely identified genes, we restricted our analysis to genes with at least one PubMed publication^46^. The final set represented 18,183 genes. Based on this set of genes, we identified their respective exons, introns, and UTRs. We defined promoter regions as the first 500 bp upstream from the transcription start site. Using our gene and DHS sets we estimated that the DHS coverage for different gene features was 26% for exons, 73% for promoters, 75% for 5'UTRs, 22% for 3'UTRs, and 20% for introns.

### Associated variants

For this study we examined variants from the GWAS catalog, variants that were eQTLs, and variants that were associated with three specific traits.

### Disease Associated Variants from the NHGRI GWAS catalog

To obtain a large set of disease-associated variants, we used the NHGRI GWAS Catalog^31^ (downloaded on November 5, 2013) and selected genome-wide significant SNPs (p-value < 5×10^−8^). To simplify the linkage disequilibrium (LD) calculations in our downstream enrichment analysis, we only included the studies that were performed in Europeans or where Europeans contributed to the majority of the final samples. We used UCSC Genome Annotations LiftOver Tool to map SNPs to the human GRCh37/hg19 reference. We excluded sex chromosomes and low frequency SNPs (MAF<0.05). For LD calculations, we used the 379 European samples from the 1000 Genomes Project^18^ available at the Beagle website (http://faculty.washington.edu/browning/beagle/beagle.html#reference_panels; Phase 1, release 2), using only bi-allelic SNPs with at least 5 copies of the minor allele. To ensure that the SNPs in our set were independent, for each pair of SNPs, we randomly excluded one SNP if r^2^ > 0.1 or if the distance between the SNPs was < 100kb. Finally, we excluded phenotypes with less than 10 independent SNP associations after our filtering criteria. This resulted in 1,416 SNPs in our test set.

### Trait-Associated Variants

To test for enrichment with trait associated variants, we used 689 SNPs reported to be associated with height^36^, 89 SNPs associated with rheumatoid arthritis in Europeans alone or shared between Europeans and Japanese^32^, and 69 SNPs associated with breast cancer^35^.

### Expression Quantitative Trait Loci

We assembled a set of cis eQTL SNPs from 923,022 SNPs associated with whole blood gene expression at a FDR of 0.5^21^. For each reported eQTL gene, we selected the single SNP most significantly associated with its expression, then performed LD pruning^47^ (using a window of 1,000 SNPs, sliding by one SNP at a time, and excluding one SNP in a pair if r^2^ > 0.1) to ensure independence of the SNPs in our final SNP set. This resulted in 6,381 SNPs in the final SNP set for whole blood gene expression.

### Simulating sets of trait-associated SNPs

To assess the performance of different enrichment tests, we wanted to define sets of SNPs that might emerge from a GWAS study where causal variants were predefined. With our approach we defined causal variants as emerging from a specific pre-defined functional region. By using this approach, we could then test the sensitivity and specificity of enrichment tests to identify appropriate enrichment.

### 1) Defining functional SNP classes

Using a total of 6,830,225 common autosomal SNPs (MAF > 5% in Europeans in the 1000 Genomes Project), we grouped SNPs into seven categories: nonsynonymous, intronic, 3' UTR, 5' UTR, promoter (<500bp from the TSS), intergenic (>5kb TSS), and those residing within DHS sites. To identify nonsynonymous variants, we used SIFT^48^. We defined promoter SNPs as the ones that mapped within 500bp upstream of the transcription start site (TSS) and intergenic SNPs as those that mapped more than 5kb upstream of the nearest TSS.

### 2) Simulating SNP sets tagging defined functional GWAS variants

We simulated 1000 sets of 1,416 causal SNPs (to match the NHGRI catalog SNP list), selected to overlap each of the predefined genomic functional categories: exons, introns, UTR's, intergenic and DHS regions. In each set, variants were randomly selected from a given functional category. For each functional variant we then identified a tagging SNP in LD that was available on a commercial genotyping array (Illumina Human Omni2.5), in order to mimic a typical GWAS approach. In total, 5,569,657 of the available SNPs were tagged (r^2^ ≥ 0.5) by 1,218,618 common (MAF ≥ 5%) variants on the Illumina Human Omni 2.5 array. If multiple SNPs were in LD with the causal variant, we selected the one with the greatest r^2^. Finally, we required SNPs in the final set to be more than 100kb apart from each other to ensure independence.

### 3) SNP sets with variable proportions of causal functional variants within DHSs.

In addition to defining SNP sets derived from causal variants within a single annotation, we also defined SNP sets where casual variants were derived from two separate functional annotations. In these instances, we selected a proportion of causal variants from one annotation, and the remainder from a second annotation.

### Inferring the proportion of NHGRI GWAS Catalog causal variants in DHS

We calculated the delta-overlap for DHS enrichment in the NHGRI GWAS Catalog, then aimed to infer the proportion of loci that overlap with DHSs from the observed delta-overlap value. We first simulated SNP sets of the same size as the GWAS Catalog, with variable proportions of causal variants within DHS, and calculated the delta-overlap in each set. We then used this relationship to infer the DHS overlap of the GWAS Catalog. Specifically, as described above, we created 1,000 SNP sets with a range of 0-45% of causal variants in DHS, in 3% increments, and calculated their corresponding delta-overlap values. (The observed delta-overlap of the real data was not consistent with any proportion that exceeded 45%,). We identified simulated sets that obtained delta-overlap values consistent with the observed delta-overlap (+/−0.2). Then, from those sets we estimated a probability distribution of proportion of causal variants within DHS. From this distribution, we estimated the mean and 95% confidence range for the proportion of causal variants in DHSs.

### Implementation

*GoShifter* is implemented as scripts written in Python-2.7. The scripts and documentation can be downloaded from www.broadinstitute.org/mpg/goshifter.

